# Hybridization, sex specific genomic architecture and local adaptation

**DOI:** 10.1101/296632

**Authors:** Anna Runemark, Fabrice Eroukhmanoff, Angela Nava-Bolaños, Jo S Hermansen, Joana I Meier

## Abstract

While gene flow can reduce the potential for local adaptation, hybridization may conversely provide genetic variation that increases the potential for local adaptation. Hybridization may also affect adaptation through altering sexual dimorphism and sexual conflict, but this remains largely unstudied. Here, we discuss how hybridization may affect sexual dimorphism and conflict due to differential effects of hybridization on males and females, and then how this in turn may affect local adaptation. First, the lower viability of the heterogametic sex in hybrids could shift the balance in sexual conflict. Second, sex-specific inheritance of the mitochondrial genome in hybrids may lead to cyto-nuclear mismatches, for example in the form of “mother’s curse”, with potential consequences for sex-ratio and sex specific expression. Third, transgressive segregation of sexually antagonistic alleles could lead to greater sexual dimorphism in hybrid populations. These mechanisms can reduce sexual conflict and enhance intersexual niche partitioning, increasing the fitness of hybrids. Adaptive introgression of alleles reducing sexual conflict or enhancing intersexual niche partitioning may facilitate local adaptation, and could favour the colonization of novel habitats. We review these consequences of hybridization on sex differences and local adaptation, and discuss how their prevalence and importance could be tested empirically.

## Introduction

Recent research has highlighted the importance of understanding sex specific local adaptation [1]. Sexual dimorphism can evolve in the same way and for the same reasons as sympatric ecological divergence and speciation [2]. Sometimes, both evolve at once [3] to maximize niche packing (see Glossary) [2-4]. In addition to classical examples such as the extreme sexual dimorphism in the beaks of the Huia [5], evidence from a wide range of taxa (e.g. birds [6], reptiles [7] and fish [8]) suggests that sexual dimorphism and niche partitioning may be important mechanisms to decrease competition for food resources between males and females. Moreover, different reproductive roles may lead to different requirements on body size, habitat use or diet. While such niche division may be advantageous, the genetic correlation between the sexes may constrain the evolution of sexual dimorphism [9]. Unless resolved, selection towards different optima may result in both sexes residing away from their fitness peaks and hence sexual conflict [9].

In spite of a long-standing research tradition investigating sex-specific viability and fitness effects of hybridization [10], and an increasing appreciation of the importance of mito-nuclear co-adaptation for hybridizing taxa [11], the effects of these phenomena on the potential for local adaptation following hybridization remain largely unexplored. Sex specific inheritance- and recombination mechanisms could affect sexual dimorphism, interlocus sexual conflict(Glossary), sex specific expression patterns or sex-ratios in hybrids (Fig. S1), but this has never been the main focus of hybridization studies. Moreover, hybridization may reshuffle sexually antagonistic alleles leading to transgressive segregation [12], which may enhance sexual dimorphism in niche use. This could dampen intersexual competition and have important consequences for ecological niche breadth.

It is increasingly recognized that under certain conditions, hybridization may have a positive impact on local adaptation [13]. Traditionally, plant ecologist viewed hybridization as potentially beneficial to adaptive evolution [14,15], while zoologists viewed it mostly as a cause of maladaptive break-down of isolating mechanisms [16]. Recent studies suggest that the tree of life is rather a net of life with frequent introgression events [13,17-19]. Currently, a plethora of examples of evolutionary consequences of hybridization, ranging from local extinction to speciation are described [13]. While adaptation to novel niches by hybrid species which have trait values that differ from those of both parent species is documented [20,21], other consequences of hybridization for local adaptation are less understood [22]. In particular, we argue that there is a gap between the multitude of studies documenting sex specific viability, sex specific expression and sex biased introgression and the lack of studies of how these factors affect sexual dimorphism in ecological niche and local adaptation in hybrid species and introgressed taxa. Here, we review how hybridization interacts with sex specific inheritance and recombination mechanisms, their effects on hybrid fitness, sex specific fitness, sex ratio and how this can lead to sexual dimorphism and/or alter the prospects for local adaptation (Fig. S1). We identify exciting areas for future research and suggest analyses to elucidate effects of hybridization on the prospects of local adaptation.

## 1) How hybridization can affect sexual conflict, sex ratio and sexual dimorphism

### 1.1) Through interactions with sex chromosomes

Almost a century ago J. B. S. Haldane [10] noted that “when in the F_1_ offspring of two different animal races one sex is absent, rare, or sterile, that sex is the heterozygous sex” (Haldane’s rule; Glossary). A closely related observation is the so-called “large X(Z) effect”(Glossary), pertaining to the disproportionate contribution of the X/Z-chromosome in causing the reduced fitness of heterogametic hybrids [23]. The principal cause of both patterns is thought to be recessive alleles with deleterious effects in hybrids having a stronger impact on the heterogametic relative to the homogametic sex, due to hemizygous expression [24]. Haldane’s rule has shown to be close to universal in both XY and ZW systems, and heteromorphic sex chromosomes show reduced introgression on the X in XY (in mammals [25], flies [26]) and the Z in ZW systems (Lepidoptera [27]; birds [28,29]).

While “Haldane’s rule” and the “large X(Z) effect” both consider alleles with the same fitness effects in males and females, sex chromosomes are expected to accumulate disproportionate numbers of sexually antagonistic alleles. This follows from their sexually asymmetric inheritance resulting in the relative effect of male- and female-specific selection acting on the sex chromosomes becoming unbalanced [30]. Dominant alleles coding for sexually antagonistic traits that benefit the homogametic sex are expected to accumulate on the X chromosome in XY systems (female-benefitting alleles) and on the Z chromosome in ZW systems (male-benefitting alleles) because they spend two-thirds of their time in the homogametic sex, while recessive alleles that favour the heterogametic sex are expected to accumulate on the Z chromosome in ZW systems and on the X chromosome in XY systems because they are rarely exposed to antagonistic selection in the homogametic sex. Modifiers that lead to reduced gene expression in the sex with lower fitness or increased expression in the sex with higher fitness are expected to subsequently evolve and accumulate [31,32].

While these properties and patterns of sex chromosome evolution have been extensively reviewed elsewhere [30,32,33], their implications for sex-specific local adaptation in hybrid populations remain poorly understood. The lower viability of the heterogametic sex may lead to biased sex-ratios in hybrid populations both in laboratory settings [23] and in wild hybrids [34]. Sex-linked gene regulation may become disrupted in hybrids resulting in abnormal gene expression. Male sterility due to disrupted sex-linked gene regulation has been observed in e.g. *Drosophila* [35,36] and hybrids between *Mus musculus* and *M. domesticus* [37]. This may potentially cause sex-specific sterility, inviability or phenotypic differences influencing sexual dimorphism.

In many taxa genetic sex determination is highly liable and differs even between closely related species (e.g. fishes [38], [39], geckos [40], and *Drosophila* [41]). Hybridization between species with different sex determining regions will result in biased sex ratios and modify interactions between sex determination and sexually antagonistic alleles. Selection against the biased sex ratio may lead to turnover of sex determination genes [42]. If a sex determination or modifier gene of one species is more closely linked to sexually antagonistic genes than the sex determiners of the other species, it may readily introgress into the other species as a result of reduced sexual conflict [43]. Sexually antagonistic alleles linked to the sex determiner may introgress in concert increasing the fitness in hybrids of both sexes. For example, in guppies [44] and cichlids [39] sex chromosome turnover has been shown to occur through introgression of sex determination genes linked to sexually-antagonistic colour pattern genes.

### 1.2) Through cytonuclear incompatibilities

The mitochondrial genome encodes specific components of the oxidative phosphorylation system used for aerobic respiration [45], and there is hence strong selection for compatibility between the mitochondria and the nuclear genome [11]. The mitochondrial genome is transmitted through the maternal lineage in most species [46]. Consequently, a male-female asymmetry in the fitness effects of mitochondrial mutations can arise [47] as mtDNA mutations that affect only males detrimentally will not be subject to natural selection. The resulting accumulation of mutations that are disadvantageous to males but benign to females is coined “mother’s curse”(Glossary) [48]. This is supported by evidence for cytoplasmic variants beneficial to females being disadvantageous to males [49,50] due to mtDNA mutations with male-biased fitness costs e.g. [47,51,52]. However, compensatory nuclear adaptations may evolve after a lag time [53]. Negative effects associated with disruption of co-evolved mito-nuclear complexes e.g. on ageing [52,54] and fertility [51,54] support the existence of such compensatory genetic variants. Cytonuclear incompatibilities arising from hybridization between diverged taxa are found in a range of taxa, e.g. birds [55,56] [57], carnivorous mice [58] flat worms [59] and plants [60,61]. Suboptimal respiration is one of the fitness costs to hybrids in flycatchers [56], carnivorous mice [58], voles [62], and chickadees [63], likely due to mito-nuclear incompatibilities. The findings of heteroplasmy in hybrids across a wide range of taxa, including mussels [64], wheat [65], birds [55,57] and *Drosophila* [66] could potentially be due to selection for paternal leakage to counteract negative fitness effects of matrilinearily inherited mitochondria [67].

Interactions between mtDNA and nDNA can lead to sex-specific global transcript responses [68]. Sex specific expression alterations could either increase or decrease sexual dimorphism, contingent on whether the expression patterns of individuals with foreign mitochondria are more similar among sexes or not. Finally, introgression of heterospecific mitochondrial variants could also have direct positive effects on population fitness through replacing mutationally loaded genomes (e.g. due to Muller’s ratchet [69]; see Glossary) as suggested in [70] and through introgression of mitochondria with allelic variants that are well adapted to e.g. the local climate c.f. [71].

Cyto-nuclear incompatibilities are also found in plants where chloroplast driven incompatibilities cause reduced hybrid fitness [72,73], which can be remedied by bi-parental chloroplast inheritance [74].

### 1.3) Through unidirectional hybridization, meiotic drive and sex-biased recombination rates

Rates of introgression may also differ between the sexes due to interspecific differences in mate preferences [75]. Additionally, sex-biased dispersal [76] may lead to increased hybridization in the dispersive sex. Unidirectional hybridization may thus contribute to differential introgression between sex-linked genes and bi-parentally inherited genes [77]. Reduced or no recombination in sex-linked markers may additionally alter their introgression rates compared to other genomic regions. In the absence of recombination, the combined effects of selection against introgression on multiple loci will lead to purging of entire introgressed chromosomes and as beneficial alleles cannot recombine away from incompatibilities, they cannot introgress [78]. Differential introgression of sex-linked genes and nuclear genes may alter sexual conflict. In many species recombination rates differ drastically between the sexes also at nuclear chromosomes, whereby in one sex recombination is either completely absent (achiasmy, e.g. Drosophila, Bombyx, Gammarus; see Glossary) or restricted to telomers (heterochiasmy, e.g. some frogs, many fishes [79]; see Glossary). In these species, alleles that are beneficial mostly to the non-recombining sex cannot introgress as easily as alleles beneficial to the recombining sex thus potentially shifting the balance of sexual conflict. Finally, meiotic drive (Glossary) can manipulate the meiotic process to distort the allelic segregation away from expected Mendelian ratios [80]. The resulting reduced fecundity favors the evolution of drive suppressors [81], and the breaking-up of these associations may affect hybrid fertility and viability [80].

### 1.4) Through transgressive sorting of sexually antagonistic variation

Hybridization may reshuffle sexually antagonistic alleles [12], leading to transgressive segregation(Glossary) of phenotypic sex-differences. This may, in turn, generate early generation hybrid populations with extreme sexual dimorphism (Fig. 1A). When sexually antagonistic alleles are fixed at different loci in the hybridizing species, hybrids could either eliminate all sources of sexual antagonism or fix sexually antagonistic alleles at several loci through recombination. The latter scenario could enable hybridizing species to evolve stronger sexuall dimorphism. Sexual dimorphism may in turn increase the carrying capacity of hybrid populations through intersexual niche partitioning [82], and may even allow hybrid species to colonize habitats that are unsuitable for their parent species. Such transgression in terms of ecological niche is well-documented in hybrid species [20,83] but it has yet to be investigated from a sexual dimorphism perspective. Strongly sexually dimorphic hybrid lineages may also be able to adapt to environments with otherwise constraining levels of sex-specific selection. For instance, [84] found that the extent of sexual size dimorphism varied across a crow hybrid zone. Moreover, the sexual dimorphism was significantly correlated both to sex-specific selection on males and altitude [84].

**Fig. 1.**
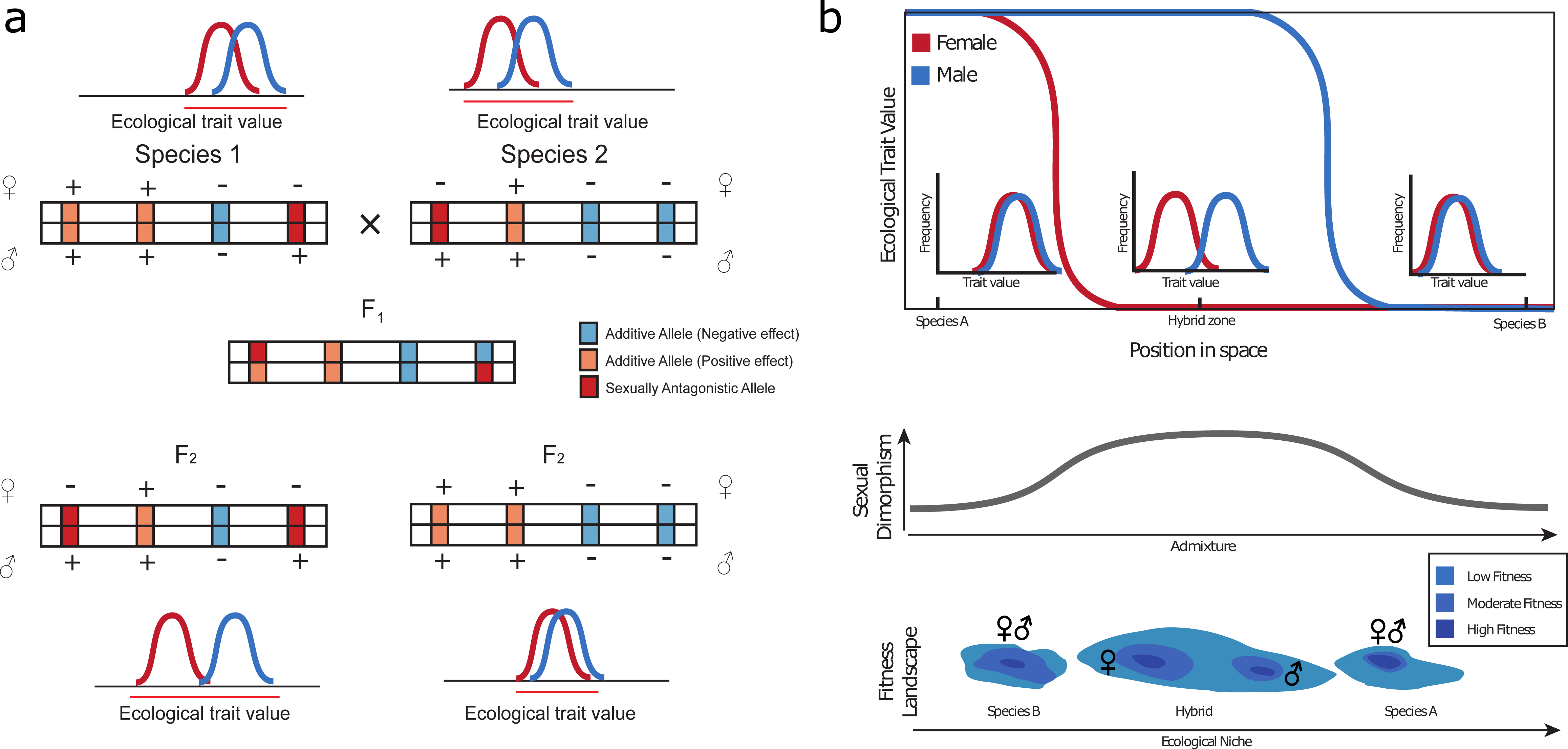
Mechanisms through which hybridization can enhance or reduce sexual dimorphism and in turn affect local adaptation. A) Transgressive segregation of sexually antagonistic alleles which have become fixed at different loci in two hybridizing species. These loci are QTL for a trait involved in niche use (e.g. beak shape in birds). After initial hybridization, recombination may lead to different phenotypic outcomes (females above and males below each locus) where sexual dimorphism is either enhanced (left lower panel) or dampened (right lower panel). This may in turn have consequences on intersexual niche partitioning and local adaptation.B) Non-coincident geographic clines between sexes for ecological traits in a hybrid zone. In admixed populations, enhanced sexual dimorphism, due to sex-specific differences in geographical clines (upper panel), may promote the occupation of novel ecological niches. Parent species may be incapable of colonizing this novel ecological niche, not because of morphospace constraints, but simply as a result of decreased mean population fitness due to intersexual competition and costly gender load (lower panel).

## 2) How hybridization may affect local adaptation via alteration of sexual dimorphism, sex ratio, and sexual conflict

### 2.1) Effects of hybridization-altered sexual dimorphism on local adaptation

As explained above, hybridization may affect sexual dimorphism via transgressive segregation of sexually antagonistic alleles or sex specific modification of expression patterns. Additionally, we argue that hybridization may affect the genetic architecture of traits in such a way that hybrid males and females reach their maximum intrinsic fitness at different levels of genome-wide admixture (for instance due to cyto-nuclear and/or sex-linked genetic incompatibilities). In hybrid zones, this may be reflected by non-coincident genomic clines(Glossary) for sex-specific genetic markers [85]. Along the hybrid zone, geographical clines of ecological traits may thus also become decoupled and displaced between males and females (Fig. 1B), especially if sex-biased genotype by environment interactions are directly affected by hybridization [86]. This could lead to a situation where sexual dimorphism increases in the centre of the hybrid zone, enhancing intersexual niche partitioning(Glossary) and mean population fitness. For two species with weak sexual dimorphism and high gender load (Glossary), hybridization could thus potentially dampen sexual conflict through formation of hybrid lineages where sexual conflict is partially or fully resolved. This would result in elevated mean population fitness, and could potentially allow for the colonization of harsh habitats where parent species would not be able to survive, c.f. [87]. Increased sexual dimorphism allows a population to explore a wider phenotypic space around the local fitness peak, potentially facilitating climbing alternative fitness peaks [88].

Finally, the impact of hybridization on sexual dimorphism could be directly involved in range shift processes(Glossary) and species range dynamics. Theory predicts that sex-specific maladaptation should increase at range margins [1]. The probability for hybridization might also increase at range margins though. Fitness asymmetries between sexes and maladaptation could thus be reduced following interspecific gene flow, and improve the viability of range margin populations [87].

### 2.2) Sex ratio distortion

Sex specific viability may result in skewed sex ratios. The Operational sex ratio (OSR; Glossary) affects the mating competition of males and females in a population [89]. The empirical evidence for this pattern is mixed [90], with one confounding factor arising as skewed sex ratios might increase the cost of mate guarding [91]. A recent meta-analysis concluded that there is compelling evidence that OSR predicts strength of sexual selection in males, but not females [92]. Sexual selection can both promote and inhibit local adaptation (reviewed in [93]). When sexual selection inhibits local adaptation, e.g. through pushing the population off the fitness optimum [94,95] a relaxation in sexual selection is likely to increase the prospects for local adaptation.

Sex ratio is also important for the ability of populations to survive and adapt as the number of females in the population determine the reproductive output e.g. [96] and strongly biased sex ratios may lead to inbreeding depression e.g. [97].

### 2.3) Effects of hybridization on local adaptation via modulation of sexual conflict

A shift in the balance between male harming and female harming antagonistic variants can lead to sex ratio distortion, which may impact local adaptation, as outlined above. In addition, a reduction of sexual conflict, e.g. due to introgression of a sex modifier increasing sex-linkage of a sexually antagonistic gene [42] or of a sex chromosome harbouring sexually antagonistic genes [87] may facilitate local adaptation by allowing for greater sexual dimorphism in ecology.

## 3) Testing for effects of hybridization on sex-specific local adaptation

Sex specific viability in early generation hybrids may result from the greater impact of deleterious recessive alleles on hybrids of the heterogametic sex, the faster X/Z theory and mitonuclear incompatibilities and lead to a biased sex ratio affecting sexual conflict and sex-specific adaptation as outlined above. Meta-analyses of sex ratios in young hybrid populations or in hybrid zones would allow to test this hypothesis, especially given such data must have been already collected and should be available from the numerous field studies of hybrid zones published over the years. Another interesting comparison would be one of sex ratios between young hybrid taxa or hybrid swarms and old, stabilized hybrid taxa. Comparing effective population sizes of the two heteromorphic sex chromosomes in hybrid taxa and parental taxa could also be informative of past sex-specific survival. Sex specific viability may affect local adaptation through relaxing sexual selection, and through increasing the probability of population persistence through female skewed sex ratios (see above). To address whether these mechanisms take place in hybrid populations, estimating the relative strength of sexual selection in hybrid taxa or hybrid zones and compare that to the parental taxa is one possibility.

Several specific predictions can be made based on the current knowledge of mitonuclear incompatibilities. First, hybrids with foreign mitochondria are expected to have suboptimal respiration and a higher incidence of sterility. Moreover, when hybrid populations differ in parental contributions (c.f. [57]) the populations with larger parts of their genomes matching the mitochondria are expected to have a more well-functioning respiration. In addition, males are expected to be disproportionately affected by mitonuclear incompatibilities. These predictions can be tested through comparing e.g. cost of respiration or basal metabolic rate in the two sexes in young hybrid taxa and stabilized hybrid taxa [62]. Moreover, meta studies addressing whether taxa with heterospecific introgressed mitochondria have obtained these from taxa adapted to the climate in their current distribution (c.f. [71]) could be interesting.

The consequences of hybridization on sexual dimorphism and local adaptation have been poorly studied, as much empirical work on hybridization often only consider one sex e.g. [98] or control for sexual dimorphism at the phenotypic level e.g. [84] without making it a specific focus. However, we argue that our hypotheses warrant reanalyses of the data on hybrid zones and hybrid species. To understand how hybridization also affects sexual dimorphism in ecological traits and niche partitioning, we suggest a more systematic investigation of whether sexual dimorphism is greater in hybrid species than in parent species. This would be predicted if transgressive sorting of sexually antagonistic alleles would enable increased dimorphism. Consistent testing of variation in sexual dimorphism across hybrid zones would also shed light on the effects of hybridization on sexual dimorphism. Another interesting possibility is to use hybrid zones as natural experiments, and test if genomic clines and geographical clines differ between sexes.

If the hybrid zone clines of ecological traits are shifted between the sexes, it implies that in the centre of the hybrid zones, ecological fitness differs between males and females (Fig. 1b). In some taxa, the clades with strongest sexual dimorphism show high rates of turnover in sex determination potentially to reduce sexual conflict and high rates of hybridization (e.g. cichlids [39] and jumping spiders [99,100]. Investigating the role of introgression in sex chromosome turnover in these systems and performing meta-analyses investigating the generality of these findings would be a promising avenue.

Little if anything is known about how the above outlined phenomena differ between early generation hybrids and stabilized hybrid taxa. Investigating this might give insights into the selection for compatibility of hybrid genomes [57,101] and the balance between this and selection for local adaptation [102]. We argue that the study of hybridization should move beyond the classical approaches and also focus on the study of ecological effects affecting sexes differently, e.g. sexual dimorphism, sex differences in viability and sexual conflict. Much remains to be done to assess the generality of the impact of hybridization on sexual genetic architecture and its consequences on adaptive potential of hybrid lineages.

## Glossary

Achiasmy: Absence of autosomal recombination in one sex.
Gender load: The reduction of fitness resulting from sexual conflict.
Genomic cline: Analysis that compares allele or genotype frequencies of each locus to a genome-wide average.
Haldane’s rule: If only one sex is inviable or sterile in a species hybrid, that sex is more likely to be the heterogametic sex.
Heterochiasmy: Differential recombination rates between sexes.
Interlocus sexual conflict: Displacement of the phenotypic optimum due to selection on the opposite sex, and by interactions between sexually antagonistic alleles at different loci.
Intersexual niche partitioning: The divergence in the niche space between the sexes.
Large X(Z) effect: Sex chromosomes (X or Z) play a disproportionate impact in adaptive evolution.
Meiotic drive: When a gene is passed to the offspring more than the expected due to manipulation of the meiotic process.
Mother’s curse: Accumulation mutations deleterious to males but not females on the mitochondria as mtDNA mutations that affect only males will not be subject to natural selection.
Muller’s ratchet: Irreversible accumulation of deleterious mutations in the genomes of asexual populations.
Niche packing: An approach to understanding the number of species and their relative abundance in dimensional niche space where the niches are packed.
Operational sex ratio (OSR): the ratio of fertilizable females to sexually active males at any given time.
Range shift processes: the processes that might shifts the range as climatic factors, dispersal capacity and population persistence.
Transgressive segregation: Progeny trait values that fall outside the range of both parents.

## Authors’ contributions

A.R. and F.E. conceived of the idea, AR wrote most of the paper and all other authors helped writing and preparing figures. All authors gave final approval for publication.

## References

1. Connallon, T. 2015 The geography of sex-specific selection, local adaptation, and sexual dimorphism. Evolution 69, 2333–2344. (doi:10.1111/evo.12737)

2. Bolnick, D. I. & Doebeli, M. 2003 Sexual dimorphism and adaptive speciation: two sides of the same ecological coin. Evolution 57, 2433–18. (doi:10.1554/02-595)

3. Cooper, I. A., Gilman, R. T. & Boughman, J. W. 2011 Sexual dimorphism and speciation on two ecological coins: Patterns from nature and theoretical predictions. Evolution 65, 2553–2571. (doi:10.1111/j.1558-5646.2011.01332.x)

4. Shine, R. 1989 Ecological causes for the evolution of sexual dimorphism: a review of the evidence. Q Rev Biol 64, 419–461. (DOI: 10.1086/416458)

5. Moorehouse, R. J. 1996 The extraordinary bill dimorphism of the Huia (*Heteralocha acutirostris):* sexual selection or intrasexual competition. Notornis 43, 19–34.

6. Radford, A. N. & Plessis, Du, M. A. 2003 Bill dimorphism and foraging niche partitioning in the green woodhoopoe. Journal of Animal Ecology 72, 258–269. (doi:10.1046/j.1365-2656.2003.00697.x)

7. Furtado, M. F. D., Travaglia-Cardoso, S. R. & Rocha, M. M. T. 2006 Sexual dimorphism in venom of Bothrops jararaca (Serpentes: Viperidae). Toxicon 48, 401–410. (doi:10.1016/j.toxicon.2006.06.005)

8. Sakashita, H. 1992 Sexual dimorphism and food habits of the clingfish, Diademichthys lineatus, and its dependence on host sea urchin. Environmental Biology of Fishes 34, 95–101. (doi:10.1007/BF00004787)

9. Rice, W. R. & Chippindale, A. K. 2008 Intersexual ontogenetic conflict. J. Evol. Biol. 14, 685–693. (doi:10.1046/j.1420-9101.2001.00319.x)

10. Haldane, J. B. S. 1922 Sex ratio and unisexual sterility in hybrid animals. Journal of Genetics 12, 101–109. (doi:10.1007/BF02983075)

11. Hill, G. E. 2016 Mitonuclear coevolution as the genesis of speciation and the mitochondrial DNA barcode gap. Ecol Evol 6, 5831–5842. (doi:10.1002/ece3.2338)

12. Parsons, K. J., Son, Y. H. & Albertson, R. C. 2011 Hybridization Promotes Evolvability in African Cichlids: Connections Between Transgressive Segregation and Phenotypic Integration. Evol Biol 38, 306–315. (doi:10.1007/s11692-011-9126-7)

13. Abbott, R. et al. 2013 Hybridization and speciation. J. Evol. Biol. 26, 229–246. (doi:10.1111/j.1420-9101.2012.02599.x)

14. Andersson, E. & Stebbins, G. L. 1954 Hybridization as an evolutionary stimulus. Evolution 4, 378–388.(http://www.jstor.org/stable/2405784)

15. Anderson, E. 1949 Introgressive hybridization. New York,: J. Wiley. (doi:10.5962/bhl.title.4553)

16. Mayr, E. 1963 Animal Species and Evolution. Cambridge, MA and London, England: Harvard University Press. (doi:10.4159/harvard.9780674865327)

17. Pennisi, E. 2016 Shaking up the tree of life. Science 354, 817–821. (DOI: 10.1126/science.354.6314.817)

18. Mallet, J. 2005 Hybridization as an invasion of the genome. Trends in Ecology & Evolution 20, 229–237. (doi:10.1016/j.tree.2005.02.010)

19. Mallet, J., Besansky, N. & Hahn, M. W. 2015 How reticulated are species? BioEssays 38, 140–149. (doi:10.1002/bies.201500149)

20. Rieseberg, L. H. 2003 Major Ecological Transitions in Wild Sunflowers Facilitated by Hybridization. Science 301, 1211–1216. (doi:10.1126/science.1086949)

21. Kagawa, K. & Takimoto, G. 2017 Hybridization can promote adaptive radiation by means of transgressive segregation. Ecol Letters 21, 264–274. (doi:10.1111/ele.12891)

22. Arnold, B. J., Lahner, B., DaCosta, J. M., Weisman, C. M., Hollister, J. D., Salt, D. E., Bomblies, K. & Yant, L. 2016 Borrowed alleles and convergence in serpentine adaptation. Proc. Natl. Acad. Sci. U.S.A. 113, 8320–8325. (doi:10.1073/pnas.1600405113)

23. Coyne, J. A. & Orr, H. A. 1989 Patterns of Speciation in Drosophila. Evolution 43, 362. (doi:10.2307/2409213)

24. Turelli, M. & Orr, H. A. 1995 The dominance theory of Haldane’s rule. Genetics 140, 389–402.

25. Janousek, V. et al. 2012 Genome-wide architecture of reproductive isolation in a naturally occurring hybrid zone between Mus musculus musculusand M. m. domesticus. Mol. Ecol. 21, 3032–3047. (doi:10.1111/j.1365-294X.2012.05583.x)

26. Garrigan, D., Kingan, S. B., Geneva, A. J., Andolfatto, P., Clark, A. G., Thornton, K. R. & Presgraves, D. C. 2012 Genome sequencing reveals complex speciation in the Drosophila simulans clade. Genome Research 22, 1499–1511. (doi:10.1101/gr.130922.111)

27. Martin, S. H. et al. 2013 Genome-wide evidence for speciation with gene flow in Heliconius butterflies. Genome Research 23, 1817–1828. (doi:10.1101/gr.159426.113)

28. Saetre, G. P., Borge, T., Lindroos, K., Haavie, J., Sheldon, B. C., Primmer, C. & Syvanen, A. C. 2003 Sex chromosome evolution and speciation in Ficedula flycatchers. Proceedings of the Royal Society B: Biological Sciences 270, 53–59. (doi:10.1098/rspb.2002.2204)

29. Ellegren, H. et al. 2012 The genomic landscape of species divergence in Ficedula flycatchers. Nature 491, 756–760. (doi:10.1038/nature11584)

30. Irwin, D. E. 2018 Sex chromosomes and speciation in birds and other ZW systems. Mol. Ecol. (doi:10.1111/mec.14537)

31. Rice, W. R. 1984 Sex chromosomes and the evolution of sexual dimorphism. Evolution 38, 735–742. (doi:10.1111/j.1558-5646.1984.tb00346.x)

32. Qvarnström, A. & Bailey, R. I. 2008 Speciation through evolution of sex-linked genes. Heredity 102, 4–15. (doi:10.1038/hdy.2008.93)

33. Schilthuizen, M., Giesbers, M. C. W. G. & Beukeboom, L. W. 2011 Haldane’s rule in the 21st century. Heredity 107, 95–102. (doi:10.1038/hdy.2010.170)

34. Veen, T., Borge, T., Griffith, S. C., Saetre, G. P., Bures, S., Gustafsson, L. & Sheldon, B. C. 2001 Hybridization and adaptive mate choice in flycatchers. Nature 411, 45–50. (doi:10.1038/35075000)

35. Michalak, P. & Noor, M. A. F. 2003 Genome-wide patterns of expression in Drosophila pure species and hybrid males. Molecular Biology and Evolution 20, 1070–1076. (doi:10.1093/molbev/msg119)

36. Moehring, A. J., Teeter, K. C. & Noor, M. A. F. 2006 Genome-Wide Patterns of Expression in Drosophila Pure Species and Hybrid Males. II. Examination of Multiple-Species Hybridizations, Platforms, and Life Cycle Stages. Molecular Biology and Evolution 24, 137–145. (doi:10.1093/molbev/msl142)

37. Good, J. M., Giger, T., Dean, M. D. & Nachman, M. W. 2010 Widespread Over-Expression of the X Chromosome in Sterile F1 Hybrid Mice. PLoS Genet 6, e1001148. (doi:10.1371/journal.pgen.1001148)

38. Woram, R. A. et al. 2003 Comparative genome analysis of the primary sex-determining locus in salmonid fishes. Genome Research 13, 272–280. (doi:10.1101/gr.578503)

39. Seehausen, O. van Alphen J.J.M. & Lande, R. 1999 Color polymorphism and sex ratio distortion in a cichlid fish as an incipient stage in sympatric speciation by sexual selection. Ecol Letters 2, 367–378. (doi:10.1046/j.1461-0248.1999.00098.x)

40. Gamble, T., Coryell, J., Ezaz, T., Lynch, J., Scantlebury, D. P. & Zarkower, D. 2015 Restriction Site-Associated DNA Sequencing (RAD-seq) Reveals an Extraordinary Number of Transitions among Gecko Sex-Determining Systems. Molecular Biology and Evolution 32, 1296–1309. (doi:10.1093/molbev/msv023)

41. Vicoso, B. & Bachtrog, D. 2015 Numerous Transitions of Sex Chromosomes in Diptera. PLoS Biol 13, e1002078–22. (doi:10.1371/journal.pbio.1002078)

42. Vuilleumier, S., Lande, R., van Alphen, J. J. M. & Seehausen, O. 2007 Invasion and fixation of sex-reversal genes. J. Evol. Biol. 20, 913–920. (doi:10.1111/j.1420-9101.2007.01311.x)

43. Bull, J. J. & Charnov, E. L. 1977 Changes in the heterogametic mechanism of sex determination. Heredity 39, 1–14. (doi:10.1038/hdy.1977.38)

44. Volff, J. N. & Schartl, M. 2001 Variability of genetic sex determination in poeciliid fishes. Genetica 111, 101–110. (https://doi.org/10.1023/A:1013795415808)

45. Björkholm, P., Harish, A., Hagström, E., Ernst, A. M. & Andersson, S. G. E. 2015 Mitochondrial genomes are retained by selective constraints on protein targeting. Proc. Natl. Acad. Sci. U.S.A. 112, 10154–10161. (doi:10.1073/pnas.1421372112)

46. Birky, C. W. 1995 Uniparental inheritance of mitochondrial and chloroplast genes: mechanisms and evolution. Proc. Natl. Acad. Sci. U.S.A. 92, 11331–11338.

47. Beekman, M., Dowling, D. K. & Aanen, D. K. 2014 The costs of being male: are there sex-specific effects of uniparental mitochondrial inheritance? Philos. Trans. R. Soc. Lond., B, Biol. Sci. 369, 20130440–20130440. (doi:10.1098/rstb.2013.0440)

48. Gemmell, N. J., Metcalf, V. J. & Allendorf, F. W. 2004 Mother’s curse: the effect of mtDNA on individual fitness and population viability. Trends in Ecology & Evolution 19, 238–244. (doi:10.1016/j.tree.2004.02.002)

49. Rand, D. M., Fry, A. & Sheldahl, L. 2005 Nuclear-Mitochondrial Epistasis and Drosophila Aging: Introgression of Drosophila simulans mtDNA Modifies Longevity in D. melanogaster Nuclear Backgrounds. Genetics 172, 329–341. (doi:10.1534/genetics.105.046698)

50. Sackton, T. B., Haney, R. A. & Rand, D. M. 2003 Cytonuclear coadapation in Drosophila: Disruption of cytochrome C oxidase in backcross genotypes. Evolution 57, 2315–2325. (DOI: 10.1111/j.0014-3820.2003.tb00243.x)

51. Patel, M. R. et al. 2016 A mitochondrial DNA hypomorph of cytochrome oxidase specifically impairs male fertility in Drosophila melanogaster. eLife 5, 1144–27. (doi:10.7554/eLife.16923)

52. Camus, M. F., Clancy, D. J. & Dowling, D. K. 2012 Mitochondria, Maternal Inheritance, and Male Aging. Current Biology 22, 1717–1721. (doi:10.1016/j.cub.2012.07.018)

53. Connallon, T., Camus, M. F., Morrow, E. H. & Dowling, D. K. 2018 Coadaptation of mitochondrial and nuclear genes, and the cost of mother’s curse. Proceedings of the Royal Society B: Biological Sciences 285, 20172257–9. (doi:10.1098/rspb.2017.2257)

54. Camus, M. F., Wolf, J. B. W., Morrow, E. H. & Dowling, D. K. 2015 Single Nucleotides in the mtDNA Sequence Modify Mitochondrial Molecular Function and Are Associated with Sex-Specific Effects on Fertility and Aging. Current Biology 25, 2717–2722. (doi:10.1016/j.cub.2015.09.012)

55. Trier, C. N., Hermansen, J. S., Sætre, G.-P. & Bailey, R. I. 2014 Evidence for Mito-Nuclear and Sex-Linked Reproductive Barriers between the Hybrid Italian Sparrow and Its Parent Species. PLoS Genet 10, e1004075–10. (doi:10.1371/journal.pgen.1004075)

56. McFarlane, S. E., Sirkiä, P. M., Ǻlund, M. & Qvarnström, A. 2016 Hybrid Dysfunction Expressed as Elevated Metabolic Rate in Male Ficedula Flycatchers. PLoS ONE 11, e0161547–10. (doi:10.1371/journal.pone.0161547)

57. Runemark, A., Trier, C. N., Eroukhmanoff, F., Hermansen, J. S., Matschiner, M., Ravinet, M., Elgvin, T. O. & Sætre G.-P. 2018 Variation and constraints in hybrid genome formation. Nature Ecology & Evolution, 1–11. (doi:10.1038/s41559-017-0437-7)

58. Shipley, J. R., Campbell, P., Searle, J. B. & Pasch, B. 2016 Asymmetric energetic costs in reciprocal-cross hybrids between carnivorous mice (Onychomys). J. Exp. Biol. 219, 3803–3809. (doi:10.1242/jeb.148890)

59. Chang, C.-C., Rodriguez, J. & Ross, J. 2016 Mitochondrial–Nuclear Epistasis Impacts Fitness and Mitochondrial Physiology of Interpopulation Caenorhabditis briggsaeHybrids. G3 6, 209–219. (doi:10.1534/g3.115.022970)

60. Barr, C. M. & Fishman, L. 2010 The Nuclear Component of a Cytonuclear Hybrid Incompatibility in Mimulus Maps to a Cluster of Pentatricopeptide Repeat Genes. Genetics 184, 455–465. (doi:10.1534/genetics.109.108175)

61. Gaborieau, L., Brown, G. G. & Mireau, H. 2016 The Propensity of Pentatricopeptide Repeat Genes to Evolve into Restorers of Cytoplasmic Male Sterility. Front Plant Sci 7, 1816. (doi:10.3389/fpls.2016.01816)

62. Boratyński, Z., Ketola, T., Koskela, E. & Mappes, T. 2015 The Sex Specific Genetic Variation of Energetics in Bank Voles, Consequences of Introgression? Evol Biol 43, 37–47. (doi:10.1007/s11692-015-9347-2)

63. Olson, J. R., Cooper, S. J. & Swanson, D. L. 2010 The relationship of metabolic performance and distribution in black-capped and Carolina chickadees. Physiol Biochem Zool. 83, 263–275.(doi:10.1086/296632.)

64. Śmietanka, B. & Burzyński, A. 2017 Disruption of doubly uniparental inheritance of mitochondrial DNA associated with hybridization area of European Mytilus edulis and Mytilus trossulusin Norway. Mar. Biol. 164, 209. (doi:10.1007/s00227-017-3235-5)

65. Aksyonova, E., Sinyavskaya, M., Danilenko, N., Pershina, L., Nakamura, C. & Davydenko, O. 2005 Heteroplasmy and paternally oriented shift of the organellar DNA composition in barley–wheat hybrids during backcrosses with wheat parents. Genome 48, 761–769. (doi:10.1139/g05-049)

66. Dokianakis, E. & Ladoukakis, E. D. 2014 Different degree of paternal mtDNA leakage between male and female progeny in interspecific Drosophila crosses. Ecol Evol 4, 2633–2641. (doi:10.1002/ece3.1069)

67. Kuijper, B., Lane, N. & Pomiankowski, A. 2015 Can paternal leakage maintain sexually antagonistic polymorphism in the cytoplasm? J. Evol. Biol. 28, 468–480. (doi:10.1111/jeb.12582)

68. Mossman, J. A., Tross, J. G., Li, N., Wu, Z. & Rand, D. M. 2016 Mitochondrial-Nuclear Interactions Mediate Sex-Specific Transcriptional Profiles in Drosophila. Genetics 204, 613–630. (doi:10.1534/genetics.116.192328)

69. Muller, H. J. 1932 Some Genetic Aspects of Sex. The American Naturalist 66, 118–138. (doi:10.1086/280418)

70. Llopart, A., Herrig, D., Brud, E. & Stecklein, Z. 2014 Sequential adaptive introgression of the mitochondrial genome in Drosophila yakuba and Drosophila santomea. Mol. Ecol. 23, 1124–1136. (doi:10.1111/mec.12678)

71. Morales, H. E., Sunnucks, P., Joseph, L. & Pavlova, A. 2017 Perpendicular axes of differentiation generated by mitochondrial introgression. Mol. Ecol. 26, 3241–3255. (doi:10.1111/mec.14114)

72. Barnard-Kubow, K. B., So, N. & Galloway, L. F. 2016 Cytonuclear incompatibility contributes to the early stages of speciation. Evolution 70, 2752–2766. (doi:10.1111/evo.13075)

73. Zeng, Y.-F., Zhang, J.-G., Duan, A.-G. & Abuduhamiti, B. 2016 Genetic structure of Populus hybrid zone along the Irtysh River provides insight into plastid-nuclear incompatibility. Sci Rep 6, 28043. (doi:10.1038/srep28043)

74. Barnard-Kubow, K. B., McCoy, M. A. & Galloway, L. F. 2017 Biparental chloroplast inheritance leads to rescue from cytonuclear incompatibility. New Phytol. 213, 1466–1476. (doi:10.1111/nph.14222)

75. Peters, K. J., Myers, S. A., Dudaniec, R. Y., O’Connor, J. A. & Kleindorfer, S. 2017 Females drive asymmetrical introgression from rare to common species in Darwin’s tree finches. J. Evol. Biol. 30, 1940–1952. (doi:10.1111/jeb.13167)

76. Greenwood, P. J. 1980 Mating systems, philopatry and dispersal in birds and mammals. Animal Behaviour 28, 1140–1162. (doi:10.1016/S0003-3472(80)80103-5)

77. Wirtz, P. 1999 Mother species-father species: unidirectional hybridization in animals with female choice. Animal Behaviour 58, 1–12. (doi:10.1006/anbe.1999.1144)

78. Martin, S. H. & Jiggins, C. D. 2017 Interpreting the genomic landscape of introgression. Current Opinion in Genetics & Development 47, 69–74. (doi:10.1016/j.gde.2017.08.007)

79. Lenormand, T. & Dutheil, J. 2005 Recombination difference between sexes: a role for haploid selection. PLoS Biol 3, e63. (doi:10.1371/journal.pbio.0030063)

80. McDermott, S. R. & Noor, M. A. F. 2010 The role of meiotic drive in hybrid male sterility. Philos. Trans. R. Soc. Lond., B, Biol. Sci. 365, 1265–1272. (doi:10.1098/rstb.2009.0264)

81. Larracuente, A. M. & Presgraves, D. C. 2012 The selfish Segregation Distorter gene complex of Drosophila melanogaster. Genetics 192, 33–53. (doi:10.1534/genetics.112.141390)

82. Butler, M. A., Sawyer, S. A. & Losos, J. B. 2007 Sexual dimorphism and adaptive radiation in Anolis lizards. Nature 447, 202–205. (doi:10.1038/nature05774)

83. Nolte, A. W., Freyhof, J., Stemshorn, K. C. & Tautz, D. 2005 An invasive lineage of sculpins, Cottus sp. (Pisces, Teleostei) in the Rhine with new habitat adaptations has originated from hybridization between old phylogeographic groups. Proceedings of the Royal Society B: Biological Sciences 272, 2379–2387. (doi:10.1038/rspb.2005.3231)

84. Saino, N. & Bernardi, F. D. 1994 Geographic variation in size and sexual dimorphism across a hybrid zone between Carrion Crows (Corvus corone corone) and Hooded Crows (C. c. cornix). Canadian Journal of Zoology 72, 1543–1550. (doi:10.1139/z94-205)

85. Jaarola, M., Tegelstrom, H. & Fredga, K. 1997 A Contact Zone with Noncoincident Clines for Sex-Specific Markers in the Field Vole (Microtus agrestis). Evolution 51, 241. (doi:10.2307/2410977)

86. Benvenuto, C., Cheyppe-Buchmann, S., Bermond, G., Ris, N. & Fauvergue, X. 2012 Intraspecific hybridization, life history strategies and potential invasion success in a parasitoid wasp. Evolutionary Ecology 26, 1311–1329. (doi:10.1007/s10682-011-9553-z)

87. Kunte, K., Shea, C., Aardema, M. L., Scriber, J. M., Juenger, T. E., Gilbert, L. E. & Kronforst, M. R. 2011 Sex chromosome mosaicism and hybrid speciation among tiger swallowtail butterflies. PLoS Genet 7, e1002274. (doi:10.1371/journal.pgen.1002274)

88. Bonduriansky, R. 2011 Sexual selection and conflict as engines of ecological diversification. The American Naturalist 178, 729–745. (doi:10.1086/662665)

89. Emlen, S. T. & Oring, L. W. 1977 Ecology, sexual selection, and the evolution of mating systems. Science 197, 215–223. (DOI: 10.1126/science.327542)

90. Rios Moura, R. & Peixoto, P. E. C. 2013 The effect of operational sex ratio on the opportunity for sexual selection: a meta-analysis. Animal Behaviour 86, 675–683. (doi:10.1016/j.anbehav.2013.07.002)

91. Klug, H., Heuschele, J., Jennions, M. D. & Kokko, H. 2010 The mismeasurement of sexual selection. J. Evol. Biol. 23, 447–462. (doi:10.1111/j.1420-9101.2009.01921.x)

92. Janicke, T. & Morrow, E. H. 2018 Operational sex ratio predicts the opportunity and direction of sexual selection across animals. Ecol Letters 21, 384–391. (doi:10.1111/ele.12907)

93. Servedio, M. R. & Boughman, J. W. 2017 The Role of Sexual Selection in Local Adaptation and Speciation. Annu. Rev. Ecol. Evol. Syst. 48, 85–109. (doi:10.1146/annurev-ecolsys-110316-022905)

94. Kirkpatrick, M. 1982 Sexual Selection and the Evolution of Female Choice. Evolution 36, 1. (doi:10.2307/2407961)

95. Lande, R. 1981 Models of speciation by sexual selection on polygenic traits. Proc. Natl. Acad. Sci. U.S.A. 78, 3721–3725.

96. Harts, A. M. F., Schwanz, L. E. & Kokko, H. 2014 Demography can favour female-advantageous alleles. Proc. Biol. Sci. 281, 20140005–20140005. (doi:10.1098/rspb.2014.0005)

97. Higashiura, Y., Ishihara, M. & Schaefer, P. W. 1999 Sex ratio distortion and severe inbreeding depression in the gypsy moth Lymantria dispar L. in Hokkaido, Japan. Heredity 83, 290–297. (DOI 10.1038/sj.hdy.6885590)

98. Bailey, R. I., Tesaker, M. R., Trier, C. N. & Saetre, G. P. 2015 Strong selection on male plumage in a hybrid zone between a hybrid bird species and one of its parents. J. Evol. Biol. 28, 1257–1269. (doi:10.1111/jeb.12652)

99. Leduc-Robert, G. & Maddison, W. P. 2018 Phylogeny with introgression in Habronattus jumping spiders (Araneae: Salticidae). BMC Evol. Biol. 18, 24. (doi:10.1186/s12862-018-1137-x)

100. Maddison, W. P. & Leduc-Robert, G. 2013 Multiple origins of sex chromosome fusions correlated with chiasma localization in Habronattus jumping spiders (Araneae: Salticidae). Evolution 67, 2258–2272. (doi:10.1111/evo.12109)

101. Eroukhmanoff, F., Bailey, R. I., Elgvin, T. O., Hermansen, J. S., Runemark, A., Trier, C. N. & Sötre, G.-P. 2017 Resolution of conflict between parental genomes in a hybrid species.(doi:https://doi.org/10.1101/102970)

102. Runemark, A., Piñeiro, L., Eroukhmanoff, F. & Sætre, G.-P. 2018 Genomic contingencies and the potential for local adaptation in a hybrid species. The American Naturalist (in press)

